# Using a structural root system model to evaluate and improve the accuracy of root image analysis pipelines

**DOI:** 10.1101/074922

**Authors:** Guillaume Lobet, Iko T. Koevoets, Manuel Noll, Patrick E. Meyer, Pierre Tocquin, Loïc Pagès, Claire Périlleux

**Affiliations:** InBioS-PhytoSYSTEMS, University of Liege, 4000 Liège, Belgium; Institut für Bio-und Geowissenschaften: Agrosphare, Forschungszentrum Jülich, D52425 Julich, Germany; Plant Cell Biology, Swammerdam Institute for Life Sciences, University of Amsterdam, 1098 XH Amsterdam, The Netherlands; INRA, Centre d'Avignon, UR 1115 PSH, Site Agroparc, 84914 Avignon cedex 9, France

**Keywords:** image analysis, root structural model, benchmarking, image library, machine learning

## Abstract

Root system analysis is a complex task, often performed with fully automated image analysis pipelines. However, the outcome is rarely verified by ground-truth data, which might lead to underestimated biases.

We have used a root model, ArchiSimple, to create a large and diverse library of ground-truth root system images (10,000). For each image, three levels of noise were created. This library was used to evaluate the accuracy and usefulness of several image descriptors classically used in root image analysis softwares.

Our analysis highlighted that the accuracy of the different traits is strongly dependent on the quality of the images and the type, size and complexity of the root systems analysed. Our study also demonstrated that machine learning algorithms can be trained on a synthetic library to improve the estimation of several root system traits.

Overall, our analysis is a call to caution when using automatic root image analysis tools. If a thorough calibration is not performed on the dataset of interest, unexpected errors might arise, especially for large and complex root images. To facilitate such calibration, both the image library and the different codes used in the study have been made available to the community.

## 1 Introduction

Roots are of utmost importance in the life of plants and hence selection on root systems represents great promise for improving crop tolerance to biotic and abiotic stresses (as reviewed in (Koevoets et al., 2016). As such, their quantification is a challenge in many research projects. This quantification is usually twofold. The first step consists in acquiring images of the root system, either using classic imaging techniques (CCD cameras) or more specialized ones (microCT, X-Ray, fluorescence, …). The next step is to analyse the pictures to extract meaningful descriptors of the root system.

To paraphrase the famous Belgian surrealist painter, Rene Magritte: “figure 1A is not a root system”. Figure 1A is an image of a root system and that distinction is important. an image is indeed a two-dimensional representation of an object, which is usually three-dimensional. Nowadays, measurements are generally not performed on the root systems themselves, but on the images, and this raises some issues.

**Figure 1.**
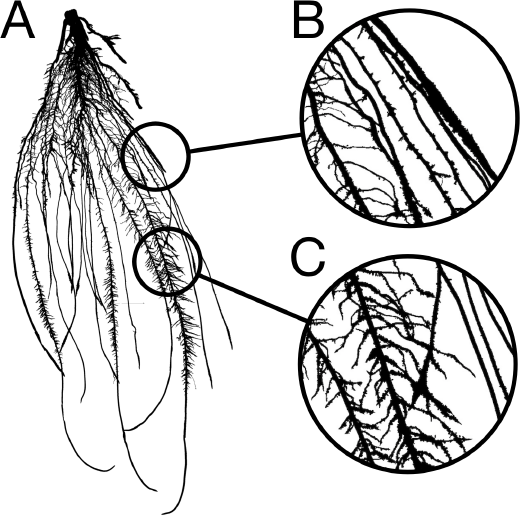
**A.** Image of a 2-week old maize root system grown in rhizotron. **B.** Close-up showing overlapping roots. **C.** Close-up showing crossing roots.

Image analysis is the acquisition of traits (or descriptors) describing the objects contained in a particular image. In a perfect situation, these descriptors would accurately represent the biological object of the image with negligible deviation from the biological truth (or data). However, in many cases, artefacts might be present in the images so that the representation of the biological object is not accurate anymore. These artefacts might be due to the conditions under which the images were taken or to the object itself. Mature root systems, for instance, are complex branched structures, composed of thousands of overlapping (fig 1B) and crossing segments (fig 1C). These features are likely to impede image analysis and create a gap between the descriptors and the data.

Root image descriptors can be separated into two main categories: morphological and geometrical descriptors. Morphological descriptors refer to the shape of the different root segments forming the root system (table 1). They include, among others, the length and diameter of the different roots. For complex root system images, morphological descriptors are difficult to obtain and are prone to error as mentioned above. Geometrical descriptors give the position of the different root segments in space. They summarize the shape of the root system as a whole. The simplest geometrical descriptors are the width and depth of the root system. Since these descriptors are mostly defined by the external envelope of the root system, crossing and overlapping segments have little impact on their estimation and hence they can be considered as relatively errorless. Geometrical descriptors are expected to be loosely linked to the actual root system topology, since identical shapes could be obtained from different root systems (the opposite is true as well). They are usually used in genetic studies, to identify genetic bases of root system shape and soil exploration.

**Table 1.**
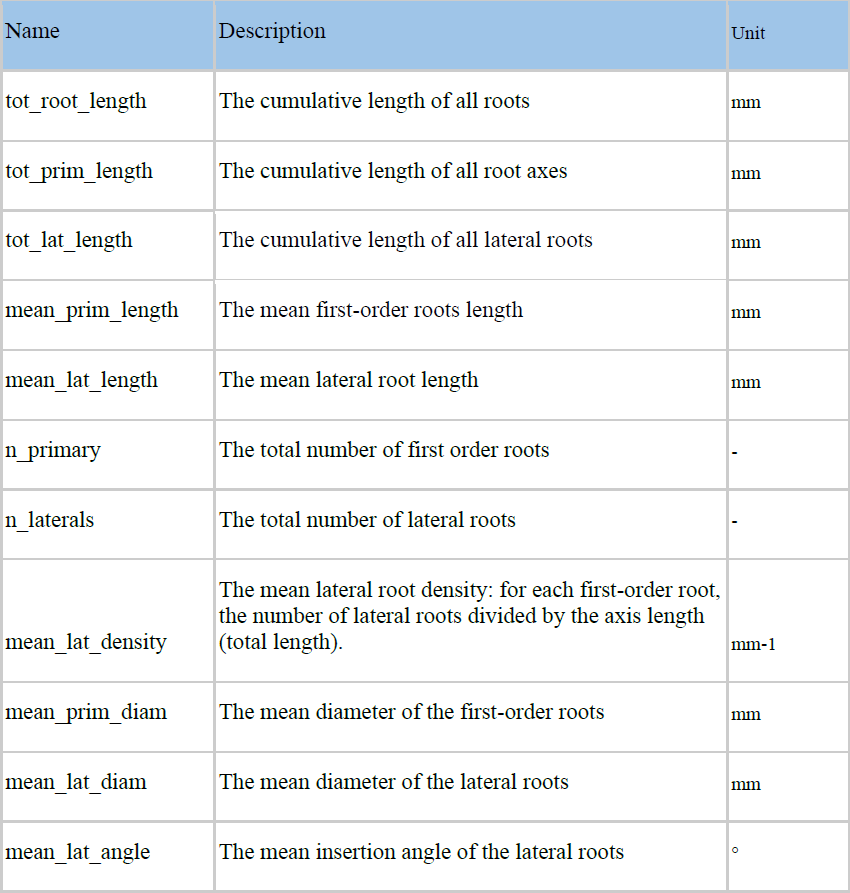
Root system parameters used as ground-truth data

Several automated analysis tools were designed in the last few years to extract both types of descriptors from root images (Armengaud et al., 2009; Bucksch et al., 2014; Galkovskyi et al., 2012; Pierret et al., 2013). However, the validation of such tools is often incomplete and/or error prone. For technical reasons, the validation is usually performed on a small number of ground-truth images of young root systems. In agreement, most analysis tools are specifically designed for this kind of root systems. In the few cases where validation is performed on large and complex root systems, it is usually not on ground-truth images, but in comparison with previously published tools (measurement of X with tool A compared with the same measurement with tool B). This might seem reasonable approach regarding the scarcity of ground-truth images of large root systems. However, the inherent limitations of these tools, such as scale or root system type (fibrous-vs. tap-roots) are often not known. Users might not even be aware that such limitations exist and apply the provided algorithm without further validation on their own images. This can lead to unexpected errors in the final measurements.

One strategy to address the lack of in-depth validation of image analysis pipelines would be to use synthetic images generated by structural root models (models designed to recreate the physical structure and shape of root systems). Many structural root models have been developed, either to model specific plant species (Pagès et al., 1989), or to be generic (Pagès et al., 2004; 2013). These models have been repeatedly shown to faithfully represent the root system structure (Pagès and Pellerin, 1996). In addition, they can provide the ground-truth data for each synthetic root system generated, independently of its complexity. However, they have not been used for validation of image analysis tools (Rellán-Álvarez et al., 2015), with one exception performed on young seedling unbranched roots (Benoit et al., 2014),.

Here we (i) illustrate the use of a structural root model, Archisimple, to systematically analyse and evaluate an image analysis pipeline and (ii) use the model-generated images to improve the estimation of root traits.

## 2 Material and methods

### 2.1 Nomenclature used in the paper

- **Ground-truth data**: The real (geometrical and morphometrical) properties of the root system as a biological object. They are determined by either manual tracking of roots or by using the output of simulated root systems.
- **(Image) Descriptor**: Property of the root image. It does not necessarily have a biological meaning.
- **Root axes**: first order roots, directly attached to the shoot
- **Lateral roots**: second-(or lower) order roots, attached to another root

### 2.2 Creation of a root system library

We used the model ArchiSimple, which was shown to allow the generation of a large diversity of root systems with a minimal amount of parameters (Pagès et al., 2013). In order to produce a large library of root systems, we ran the model 10,000 times, each time with a random set of parameters (fig. 2A). For each simulation, the growth and development of the root system were constrained in two dimensions.

The simulations were divided into two main groups: fibrous and tap-rooted. For the fibrous simulations, the model generated a random number of root axes and secondary (radial) growth was disabled. For tap-root simulations, only one root axis was produced and secondary growth was enabled (the extent of which was determined by a random parameter).

The root system created in each simulation was stored in a Root System Markup Language (RSML) file. Each RSML file was then read by the RSML Reader plugin from ImageJ to extract ground-truth data for the library (Lobet et al., 2015). These ground-truth data included geometrical and morphological parameters (table 1). For each RSML data file, the RSML Reader plugin also created three JPEG images (at a resolution of 300 DPI) for each root system, with different levels of noise (using the Salt and Pepper Filter in ImageJ) (fig 2D). For each root system, we computed overlapping index as the number of root segments having an overlap with other root segments over the total number of root segments.

### 2.3 Root image analysis

Each generated image was analysed using a custom-made ImageJ plugin, Root Image Analysis-J (or RIA-J). For each image, we extracted a set of classical root image descriptors, such as the total root length, the projected area and the number of visible root tips (fig 2E). In addition, we included shape descriptors such as the convex-hull area or the exploration ratio (see Supplemental file 1 for details of RIA-J). The list of traits and algorithms used by our pipeline is listed in table 2.

**Figure 2.**
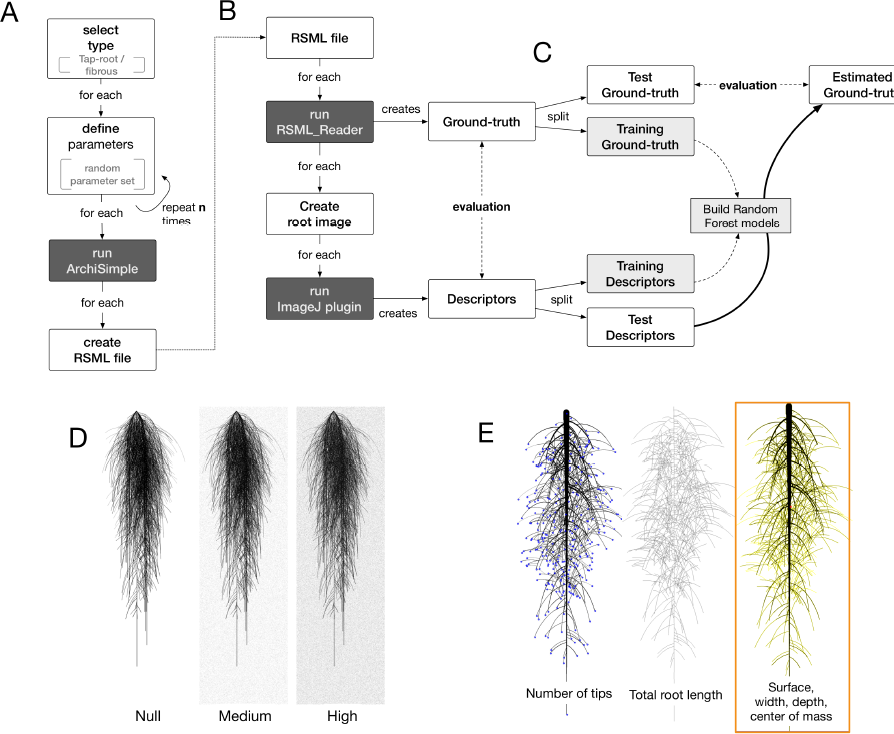
Overview of the workflow used in this study. A. Generation of root systems using Archisimple. B. Creation and analysis of root images. C. Use of Random Forest algorithms to better estimate root system ground-truths. D. Illustration of the different noise levels used in the analysis. E. Example of descriptors extracted with RIA-J.

**Table 2.** Root image descriptors extracted by RIA-J

| Name | Description | Unit | Reference |
| --- | --- | --- | --- |
| **Morphology** |  |  |  |
| area | Projected area of the root system | mm<sup>2</sup> | (Galkovskyi et al., 2012) |
| length | Length of the skeleton of the root system image | mm | (Galkovskyi et al., 2012) |
| tip_count | Number of end branches in the root system skeleton | - |  |
| diam_mean | Mean diameter of the root object in the image | mm |  |
| **Geometry** |  |  |  |
| width | The maximal width of the root system | mm | - |
| depth | The maximal depth of the root system | mm | - |
| width_depth_ratio | Ratio between the width and the depth of the root system | - | (Galkovskyi et al., 2012) |
| com_x - com_y | Relative coordinates of the centre of mass of the root system | - | (Galkovskyi et al., 2012) |
| convexhull | Area of the smallest convex shape encapsulating the root system | mm<sup>2</sup> | (Galkovskyi et al., 2012) |
| exploration | Ratio between the convex hull area and the projected area | - | (Galkovskyi et al., 2012) |

### 2.4 Data analysis

Data analysis was performed in R (R Core Team). Plots were created using *ggplot2* (Wickham, 2009) and *lattice* (Sarkar, 2008).

The Mean Relative Errors (MRE) were estimated using the equation:

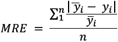
where *n* is the number of observations, 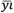 is the ground-truth and *yi* is the estimated ground-truth.

### 2.5 Random Forest Framework

A *random forest* is a state-of-the-art machine learning algorithm typically used for making new predictions (in both classification and regression tasks). Random Forests can perform non-linear predictions and, thus, those often outperform linear models. Since its introduction by Breiman in 2001 (Breiman, 2001), those have been widely used in many fields from gene regulatory network inference to generic image classification (Marée et al., 2016, Huynh-Thu et al., 2013). Random forest relies on growing a multitude of decision trees, a prediction algorithm that has shown good performances by itself but, when combined with other decision trees (hence the name forest), returns predictions that are much more robust to outliers and noisy data (see bootstrap aggregating, Breiman 1996).

In a machine learning setting one is given a set 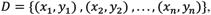 where 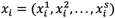 is an element of a *S* –dimensional *feature space X*, and 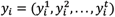 an element of a *t* –dimensional *response space Y*.

The learning task is to find a model 
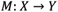
that predicts the data in a good way, where goodness is measured w.r.t. an error function *L*.

A decision tree *T*_*D*_ is a machine learning method that, for a dataset *D*, constructs a binary tree with each node representing a binary question and each leaf a value of the response space. In other words, a prediction can be made from an input value by looking at the set of binary questions that leads to a leaf (e.g. is the primary root bigger than q1 and if yes is the number of secondary roots smaller than q2 and if no, …)

Each decision is based upon exactly one feature and is used for deciding which branch of the tree a given input value must take. Hence a decision tree splits successively the set *D* into smaller subsets and assigns them a value *y*_*i*_ = *T*_*D*_(*x*_*i*_) of the response space.

The choice of the feature used for splitting depends on a relevance criterion. In our setting, the default relevance criterion from the randomForest R package (CRAN randomForest, 2015), namely the Gini index, has been used.

A random forest

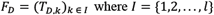

consists of *l* decision trees *T*_*Dk*_, where several key parameters such as the feature space, are chosen randomly (hence the word Random in the algorithm name). While using a random subspace strongly accelerates the growth of a single tree, it can also decrease its accuracy. However, the use of large number of trees counterbalance advantageously those two effects. The final prediction for each input value *x*_*i*_ corresponds to the majority vote of all the decision trees of the forest *T*_*D,k*_(*x*_*i*_) in a classification setting while an average of all predicted values is used in a regression task.

#### 2.5.1 Framework description

Our method consists of three typical steps:

- a preprocessing step, where we replace missing values of the training set.
- a model generation step where, for each response variable, we generate different models according to two Random Forest parameters (number of trees and number of splits).
- a model selection step, where we choose the best performing pair of parameters of the previous step for each one of the response variable.

#### 2.5.2 Preprocessing

Missing values in our dataset might arise due to highly noisy images, where the measurement of certain descriptors has been infeasible. To deal with this issue, we first replaced missing values.

This is done using the imputation function of the randomForest R package. It replaces all missing values of a response variable by the median and then a Random Forest is applied on the completed data to predict a more accurate value. We favored ten trees for computing the new value over the default value of 300 as we found that it offered sufficiently accurate results for our application while being much faster.

#### 2.5.3 Model Generation

In the model generation step, for each of the response variables, several forests with different number of trees and different number of splits (*t*_*i*_, *m*_*j*_) are tested. In practice, the training set *D*_*train*_ is divided into *m*_*j*_ disjunct subsets 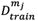 and on each of those, a random forest 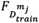 is trained on a growing number of *t*_*i*_ random trees.

#### 2.5.4 Model selection

Given a new data point *x*, each model predicts a response variable *y* by averaging the predicted values 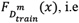, i.e.

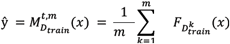

Then in a final step an estimate of the root-mean-square (RMSE) generalized error on the test set ***D*_*test*_** is computed, where RSME is defined as

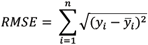

for 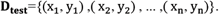

Finally, the model with the parameter pair (*t*, *m*) having the minimal error (on the separate test set) is chosen in order to make the predictions.

### 2.6 Data availability

All data used in this paper (including the image and RSML libraries) are available at the address http://doi.org/10.5281/zenodo.208214

An archived version of the codes used in this paper is available at the address http://doi.org/10.5281/zenodo.208499

An archived version of the machine learning framework is available at the address https://github.com/FaustFrankenstein/RandomForestFramework/releases/tag/v1.0

## 3 Results and discussions

### 3.1 Production of a large library of ground-truth root system images

We combined existing tools into a single pipeline to produce a large library of ground-truth root system images. The pipeline combines a root model (ArchiSimple (Pagès et al., 2013)), the Root System Markup Language (RSML) and the RSML Reader plugin from ImageJ (Lobet et al., 2015). In short, ArchiSimple was used to create a large number of root systems, based on random input parameter sets. Each output was stored as an RSML file (fig. 2A), which was then used by the RSML Reader plugin to create a graphical representation of the root system (as a.jpeg file) and a ground-truth dataset (fig. 2B). Details about the different steps are presented in the Materials and Methods section.

We used the pipeline to create a library of 10,000 root system images, separated into fibrous (multiple first order roots and no secondary growth) and tap-root systems (one first order root and secondary growth). The ranges of the different ground-truth data are shown in table 3 and their distribution is shown in the Supplemental Figure 1.

We started by evaluating whether fibrous and tap-root systems should be separated during the analysis. We performed a Principal Component Analysis on the ground-truth dataset to reduce its dimensionality and assess if the *type* grouping influenced the overall dataset structure (fig. 3A). Fibrous and tap-root systems formed distinct groups (MANOVA p-value < 0.001), with limited overlap. The first principal component, which represented 30.9% of the variation within the dataset, was mostly influenced by the number of primary axes. The second principal component (19.1% of the variation) was influenced, in part, by the root diameters. These two effects were consistent with the clear root system type grouping, since they expressed the main difference between the two groups of root-system *types*. Therefore, since the *type* grouping had such a strong effect on the overall structure, we decided to separate them for the following analyses.

**Figure 3.**
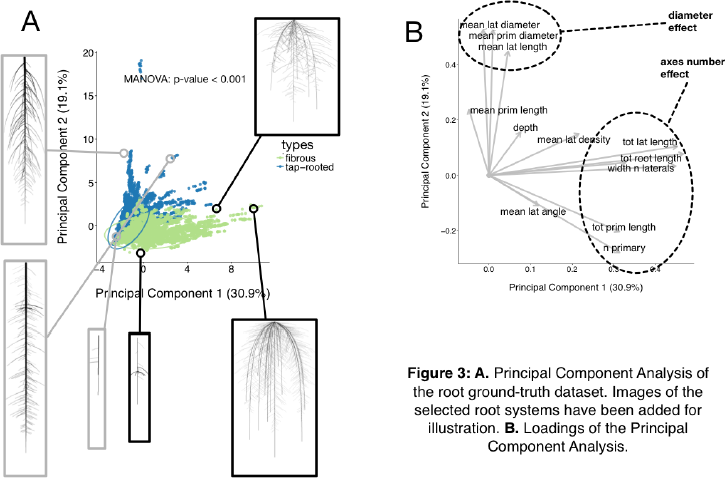
A. Principal Component Analysis of the root ground-truth dataset. Images of the selected root systems have been added for illustration. B. Loadings of the Principal Component Analysis.

**Table 3.**
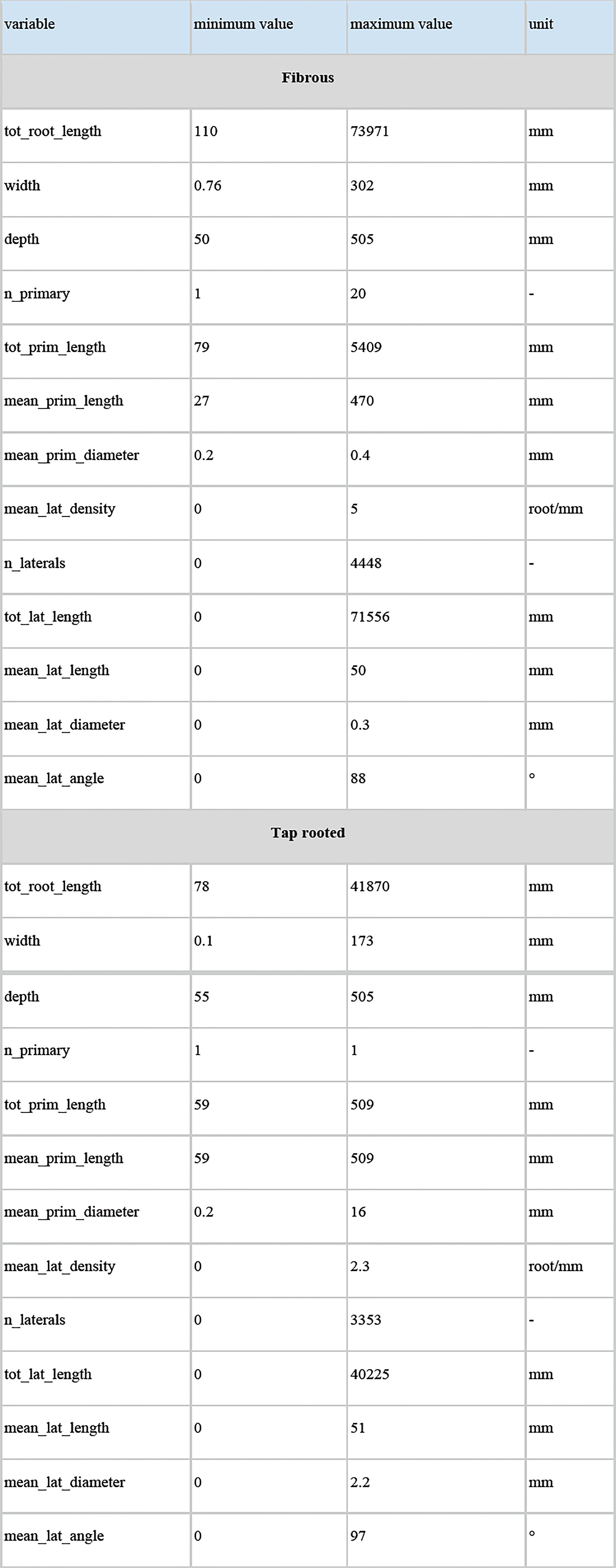
Ranges of the different ground-truth data from the root systems generated using ArchiSimple

| variable | minimum value | maximum value | unit |
| --- | --- | --- | --- |
| **Fibrous** |  |  |  |
| tot_root_length | 110 | 73971 | mm |
| width | 0.76 | 302 | mm |
| depth | 50 | 505 | mm |
| n_primary | 1 | 20 | - |
| tot_prim_length | 79 | 5409 | mm |
| mean_prim_length | 27 | 470 | mm |
| mean_prim_diameter | 0.2 | 0.4 | mm |
| mean_lat_density | 0 | 5 | root/mm |
| n_laterals | 0 | 4448 | - |
| tot_lat_length | 0 | 71556 | mm |
| mean_lat_length | 0 | 50 | mm |
| mean_lat_diameter | 0 | 0.3 | mm |
| mean_lat_angle | 0 | 88 | ° |
| **Tap rooted** |  |  |  |
| tot_root_length | 78 | 41870 | mm |
| width | 0.1 | 173 | mm |
| depth | 55 | 505 | mm |
| n_primary | 1 | 1 | - |
| tot_prim_length | 59 | 509 | mm |
| mean_prim_length | 59 | 509 | mm |
| mean_prim_diameter | 0.2 | 16 | mm |
| mean_lat_density | 0 | 2.3 | root/mm |
| n_laterals | 0 | 3353 | - |
| tot_lat_length | 0 | 40225 | mm |
| mean_lat_length | 0 | 51 | mm |
| mean_lat_diameter | 0 | 2.2 | mm |
| mean_lat_angle | 0 | 97 | ° |

### 3.2 Systematic evaluation of root image descriptors

To demonstrate the utility of a synthetic library of ground-truth root systems, we analysed every image of the library using a custom-built root image analysis tool, RIA-J. We decided to do so since our purpose was to test the usefulness of the synthetic analysis and not to assess the accuracy of existing tools. Nonetheless, RIA-J was designed using known and published algorithms, often used in root system quantification. A detailed description of RIA-J can be found in the Materials and Methods section and Supplemental File 1.

We extracted 10 descriptors from each root system image (Table 2) and compared them with their own ground-truth data. For each pair of descriptor-data, we performed a linear regression and computed its r-squared value. Figure 4 shows the results from the different combinations for both root system types. We can observe that, generally, correlations were poor with only 3% of the combinations having an r-squared above 0.8. In addition, for some ground-truth data, such as the mean lateral length or the number of primary roots, none of the descriptors actually gave a good estimation (fig 4, highlighted with arrows).

**Figure 4.**
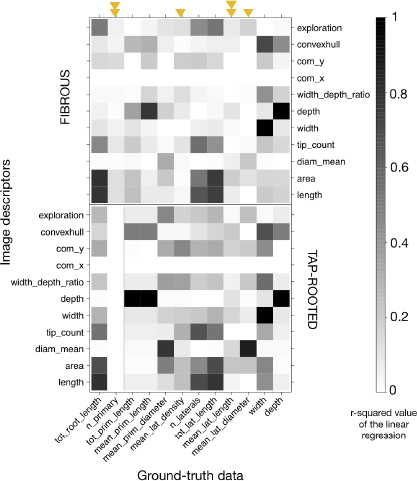
Heatmap of the r-squared values between the different image descriptors and the ground-truth values, for the images without any noise. Black represents an r-squared value of 1; white represents a value of 0. Upper panel: tap-root dataset. Lower panel: fibrous root dataset. Arrows highlight the ground-truth data that cannot be accurately described with the different descriptors. The arrows were doubled when it was the case for both fibrous and tap-rooted root systems.

Additionally, it should be noted that the correlations were different for fibrous-and tap-root systems. As an example, the correlation found between the *mean_lat_diameter* and *diam_mean* estimators was better for fibrous roots than within the tap-root dataset. Consequently, validation of the different image analysis algorithms should be performed, at least, for each group. An algorithm giving good results for a fibrous root system might fail when applied to tap-rooted ones.

### 3.3 Errors from image descriptors are likely to be non-linear across root system sizes and image qualities

In addition to being related to the species of study, estimation errors are likely to increase with the root system size. As the root system grows and develops, the number of crossing and overlapping segments increases (fig. 5A), making the subsequent image analysis potentially more difficult and prone to error. However, a systematic analysis of such error is seldom performed.

Figure 5 shows the relationship between the ground-truth and descriptor values for three parameters: the total root length (fig. 5B), the number of roots (fig. 5C) and the root system depth (fig. 5D). For each of these variables, we quantified the Mean Relative Error (see Materials and Methods for details) as a function of the overlap index. This was done for three levels of noise added to the images (“null”, “medium” and “high”). We can observe that for the estimation of both the total root length and the number of lateral roots, the Mean Relative Error increased with the size of the root system (fig. 5B–5C). As stated above, such increase of the error was somehow expected with increasing complexity. Moreover, depending on the metric of interest, such as the number of root tips, low image quality can result in high level of error. For other traits, such as the root system depth, no errors were expected (*depth* is supposedly an error-less variable) and the Mean Relative Error was close to 0 whatever the size of the root system and image quality.

The results presented here are tightly dependent on the specific algorithms used for image analysis and hence might be different for other published tools. However, they are a call for caution when analysing root images: unexpected errors in ground-truth estimation can arise. Our image library can be used to better identify the errors generated by other analysis tools, current or future.

**Figure 5.**
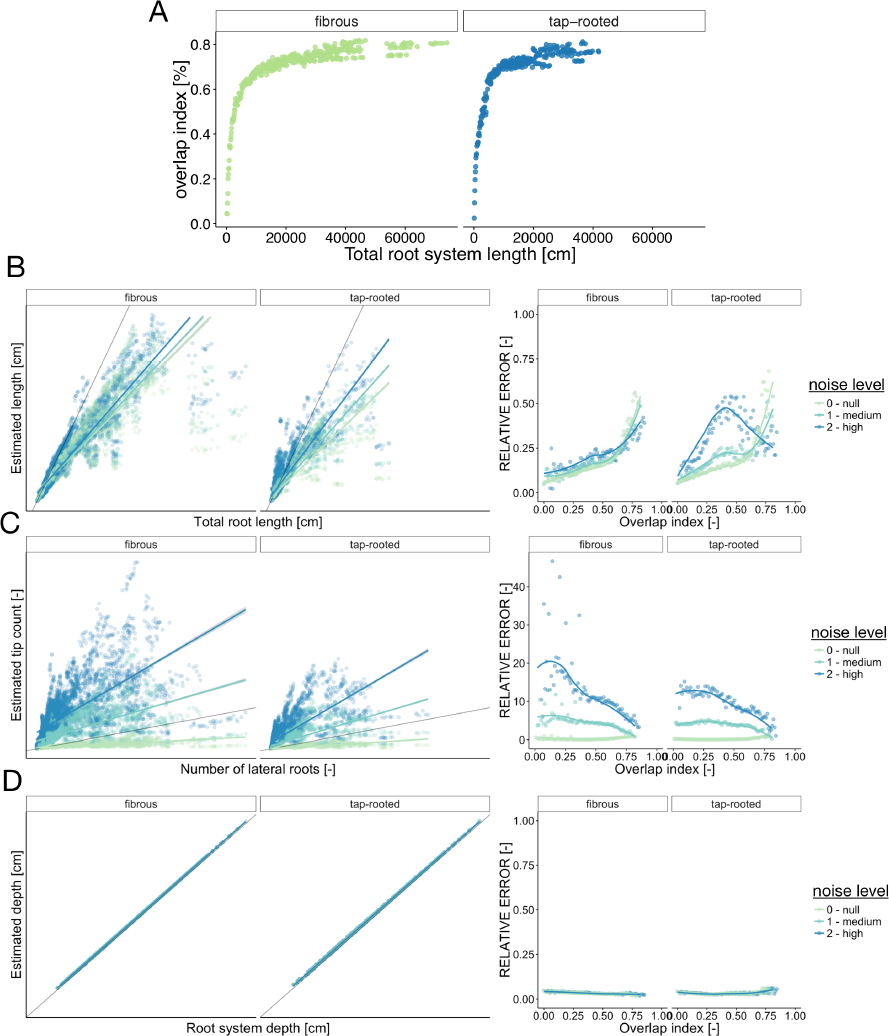
Error estimation for three ground-truth parameters. A. Evolution of the overlap index (proportion of root overlapping) with the root system size. B-D: Left panel shows the relationship between the descriptors and the corresponding ground-truth variables. Right panels show the evolution of the Mean Relative Error (MRE) as a function of the overlap index. For the MRE calculations, the continuous variables were discretized in groups. B. Total root length. C. Number of lateral roots. D. Root system depth.

### 3.4 Using the synthetic library to train machine learning algorithms

The main advantage of creating a synthetic library is to generate paired datasets of image descriptors and their corresponding ground-truth values. Having both information can, in theory, be used to either calibrate the image analysis pipeline or to identify the best descriptors for the ground-truth traits of interest. Here, we explored the second approach and used a random forest algorithm to find which combination of descriptors would best describe each ground-truth data (see Material and Methods for details). In short, we randomly divided the whole dataset into training (3/4) and testing subsets (1/4). The training set was used to create a random forest model for each ground-truth data, which was then we applied to the test set. The accuracy of these new predictions was then compared to the accuracy of the direct method (single descriptors) (fig. 2C).

Figure 6 shows the comparison of the accuracy (both the r-squared values from linear regressions and the Mean Relative Error, MRE) of both methods for each ground-truth data. We can clearly see that the random forest approach performed always better (sometimes substantially) than the direct approach, even for images with high level of noise. In addition, for most traits, the r-squared and MRE values were above 0.9 and below 0.1 respectively, which is very good, especially for such a wide range of images. In addition, the random forest approach allowed the correct estimation of traits that were difficult to estimate with the direct approach (such as the number of primary axes or the mean lateral root density).

**Figure 6.**
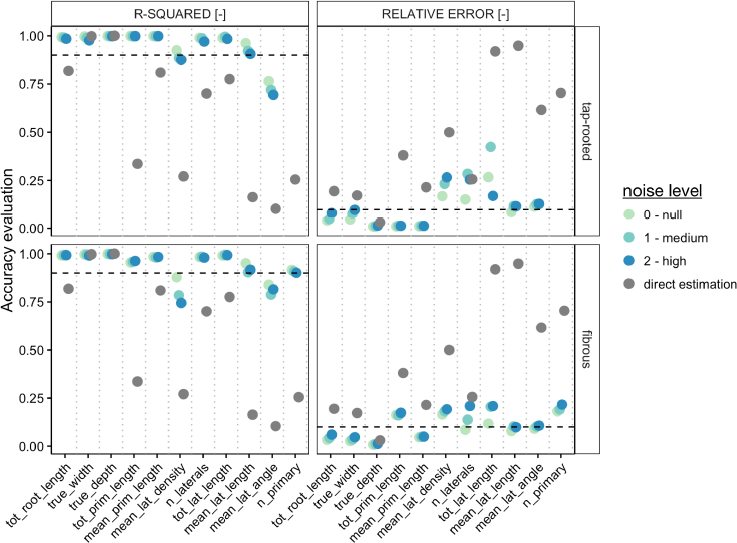
Comparison between the direct trait and the random forest approach, for the different root system types and the different levels of noise. For each metric, we computed both the r-squared value from the linear regression between the estimation and the ground-truth (left panels), as well as the Mean Relative Error (right panel). The grey points represent the values obtained with the direct estimation (best descriptor, no noise). Color points represent the values obtained with the random forest approach, for different levels of noise. The dotted lines show the 0.9 (r-squared) and 0.1(MRE) thresholds.

Figure 7 shows the detailed comparison of both methods for the estimation of the total root length. Again, a clear improvement was visible with the Random Forest method, leading to small errors, even with large root systems and noisy images.

In our study, machine learning algorithms on simulated datasets seems to yield very good results and we believe they open new avenues for root system analyses. It is clear however that their value relies on the quality and relevance of the training dataset vs. the test dataset and that they must be carefully designed.

**Figure 7.**
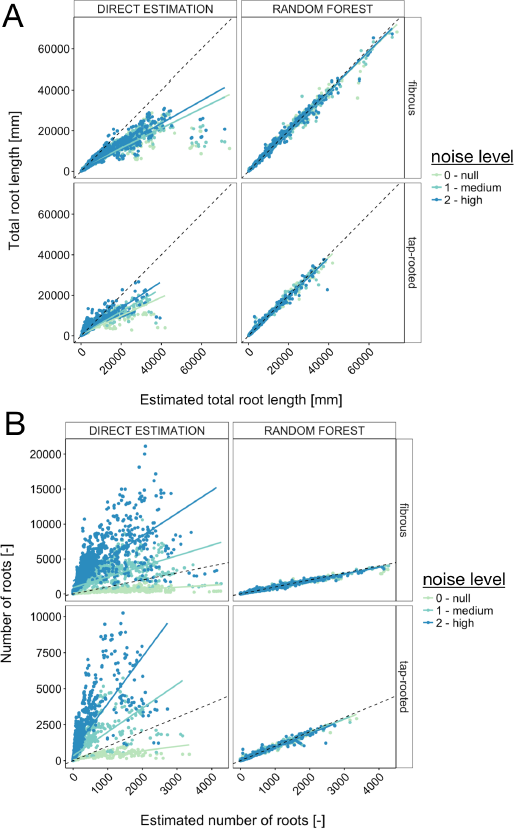
Comparison between the direct trait estimation and the random forest approach, for the different root system types and the different levels of noise. A. Comparison, for the total root length, of the accuracy of both approaches. The dotted line represents the diagonal. The plain line represents the linear regression. B. Same, for the number of roots.

## 4 Conclusions

The automated analysis of root system images is routinely performed in many research projects. Here we used a library of 10;000 synthetic images to estimate the accuracy and usefulness of different image descriptors extracted with a homemade root image analysis pipeline. Our study highlighted some limitations and biases of the image analysis process.

We found that the type of root system (fibrous vs tap-rooted), its size and complexity, as well as the quality of the images had a strong influence on the accuracy of some commonly used image descriptors and their meaning and relevance for ground-truth extraction. So far, a large proportion of the root research has been focused on seedlings with small root systems and has *de facto* avoided such errors.

However, as the research questions are likely to focus more on mature root systems in the future, these limitations will become critical. We showed that synthetic datasets can be used for calibration or modelling (machine learning) steps that allow ground-truth extraction from comparable images. We then hope that our library will be helpful for the root research community to evaluate and improve other image analysis pipelines.

## 5 Conflict of Interest

The authors declare that the research was conducted in the absence of any commercial or financial relationships that could be construed as a potential conflict of interest.

## 6 Author Contributions

GL, LP, PT, IK, MN and CP designed the study. IK developed the image analysis pipeline RIA-J. MN and PM developed the Random Forest framework. GL generated the image library and did the data analysis. LP developed the Archisimple model. All authors have participated in the writing of the manuscript.

## 7 Funding

This research was funded by the Interuniversity Attraction Poles Programme initiated by the Belgian Science Policy Office, P7/29. GL and MN are grateful to the F.R.S.-FNRS for a postdoctoral research grant (1.B.237.15F) and doctoral grant (1.A.320.16F), respectively.

## 8 Supplementary Material

**Supplemental figure 1**: Distribution of the properties of the modelled root images
**Supplemental figure 2**: Distribution of the descriptors of the modelled root images
**Supplemental file 1**: Definitions of the different descriptors extracted by RIA-J

## References

1. Armengaud, P., Zambaux, P., Hills, A., Sulpice, R., Pattison, R. J., Blatt, M. R.et al. (2009) EZ-Rhizo: integrated software for the fast and accurate measurement of root system architecture. Plant J 57, 945–956.

2. Benoit, L., Rousseau, D., Belin, É., Demilly, D., and Chapeau-Blondeau, F. (2014). Simulation of image acquisition in machine vision dedicated to seedling elongation to validate image processing root segmentation algorithms. Computers and Electronics in Agriculture 104: 84–92.

3. Bonhomme, V., Picq, S., and Gaucherel, C. (2014). Momocs: outline analysis using R. Journal of Statistical….

4. Breiman, L (2001) Random Forests. Machine Learning 45 (1): 5–32.

5. Breiman, L (1996) Bagging predictors. Machine Learning 24 (2): 123–140.

6. Bucksch, A., Burridge, J., York, L. M., Das, A., Nord, E., Weitz J. S., et al. (2014) Image-based high-throughput field phenotyping of crop roots. Plant Physiol 166: 470–486.

7. Chitwood, D. H and Otoni, W. C. (2016) Morphometric analysis of Passiflora leaves I: the relationship between landmarks of the vasculature and elliptical Fourier descriptors of the blade. *bioRxiv*.

8. CRAN randomForest (2015). https://cran.r-project.org/web/packages/randomForest/randomForest.pdf

9. Dryden, I. L. (2015). shapes package. Vienna, Austria.

10. Galkovskyi, T., Mileyko, Y., Bucksch, A., Moore, B., Symonova, O., Price, C. A., et al. (2012) GiA Roots: software for the high throughput analysis of plant root system architecture. BMC Plant Biol 12, 116.

11. Huynh-Thu V. A, Wehenkel L, & Geurts P. (2013) Gene regulatory network inference from systems genetics data using tree-based methods.

12. Koevoets, I. T., Venema, J. H., Elzenga, J. T. M., and Testerink, C. (2016) Roots Withstanding their Environment: Exploiting Root System Architecture Responses to Abiotic Stress to Improve Crop Tolerance. Front Plant Sci 07: 91–19.

13. Lobet, G., Pound, M. P., Diener, J., Pradal, C., Draye, X., Godin C., et al. (2015) Root System Markup Language: Toward a Unified Root Architecture Description Language. Plant Physiol 167: 617–627.

14. Marée, R., Geurts, P., &. Wehenkel L. (2016) Towards Generic Image Classification using Tree-based Learning: an Extensive Empirical Study. *Pattern Recognition Letters*.

15. Pagè, L., and Pellerin, S. (1996) Study of differences between vertical root maps observed in a maize crop and simulated maps obtained using a model for the three-dimensional architecture of the root system. Plant and Soil 182: 329–337.

16. Pagès, L., Bécel, C., Boukcim, H., Moreau, D., Nguyen, C., and Voisin A.-S., (2013) Calibration and evaluation of ArchiSimple, a simple model of root system architecture. Ecological Modelling 290: 76–84.

17. Pagès, L., Jordan, M. O., and Picard, D. (1989) A simulation model of the three-dimensional architecture of the maize root system. Plant and Soil 119: 147–154.

18. Pagès, L., Vercambre, G., Drouet, J.-L., Lecompte, F., Collet, C., and LeBot J. (2004) RootTyp: a generic model to depict and analyze the root system architecture. Plant and Soil 258, 103–119.

19. Pierret, A., Gonkhamdee, S., Jourdan C., and Maeght J.-L. (2013) IJ-Rhizo: an open-source software to measure scanned images of root samples. *Plant and Soil*. 1–9

20. Core R., Team, R.: A Language and Environment for Statistical Computing.

21. Rellán-Álvarez, R., Lobet, G., Lindner H., Pradier P.-L. Sebastian J. Yee M.-C. et al. (2015) GLO-Roots: an imaging platform enabling multidimensional characterization of soil-grown root systems. *PeLife* 4, e07597.

22. Ristova, D., Rosas, U., Krouk G., Ruffel S. Birnbaum K. D. Coruzzi G. M. (2013) RootScape: A landmark-based system for rapid screening of root architecture in Arabidopsis thaliana. *Plant Physiology*.

23. Sarkar, D. (2008) Lattice: Multivariate Data Visualization with R. New York: Springer.

24. Wickham, H. (2009). ggplot2. New York, NY: Springer New York

